# PACESS: Practical AI-based Cell Extraction and Spatial Statistics for large 3D bone marrow tissue images

**DOI:** 10.1101/2022.12.29.521787

**Authors:** George Adams, Floriane S. Tissot, Chang Liu, Chris Brunsdon, Ken R. Duffy, Cristina Lo Celso

## Abstract

Despite much being known about the molecular regulation of hematopoiesis, little is understood about how hematopoietic cells are organized within bone marrow (BM) tissue. Recent advances in microscopy have led to the creation of increasingly detailed images of murine hematopoietic tissue. Accurate, efficient, and informative methodologies to extract and analyze the large amount of data generated are, however, still lacking. Indeed, cells are very densely packed in the bone marrow and therefore difficult to efficiently and accurately segment. In addition, currently employed statical analyses of cellular localization are generally unsuitable for the investigation of interactions between more than two cell types. To overcome these limitations, we developed PACESS, a readily applicable method based on neural network classification of hundreds of thousands of cells in thick 3D bone marrow samples, and a combination of statistical techniques to assess the spatial interactions between multiple cell types. To validate this approach, we used it to investigate the spatial organization of T cells, megakaryocytes and leukemic cells. We demonstrate that the presence of large clusters of leukemic cells affects the distribution of both T cells and megakaryocytes, albeit differently, resulting in the generation of previously unrecognized, unique microenvironments adjacent to each other within the same bone marrow cavity. We believe that this approach can contribute to unravel the BM cellular organization.

**MOTIVATION:** The organization of diverse hematopoietic cells within bone marrow tissue remains unclear. Recently developed tissue clearing methods enable the generation of large, 3D, single cell resolution microscopy images datasets, but the bottleneck in their analysis lies in both the identification and classification of cells, and in statistical analyses to probe their spatial relationships. We present a workflow (PACESS) that takes advantage of a convoluted neural network to identify and classify cells in 2D coupled with an automated method that extrapolates to 3D, followed by a combination of spatial statistics to classify tissue regions based on each cell type’s density, and logistic regression to test whether the relative abundance of cell types may be explained by reciprocal dependencies. Finally, we provide a combined measurement of the abundance of all cell types in a 3D map, highlighting regional variations found within the tissue.

**Graphical Abstract:** 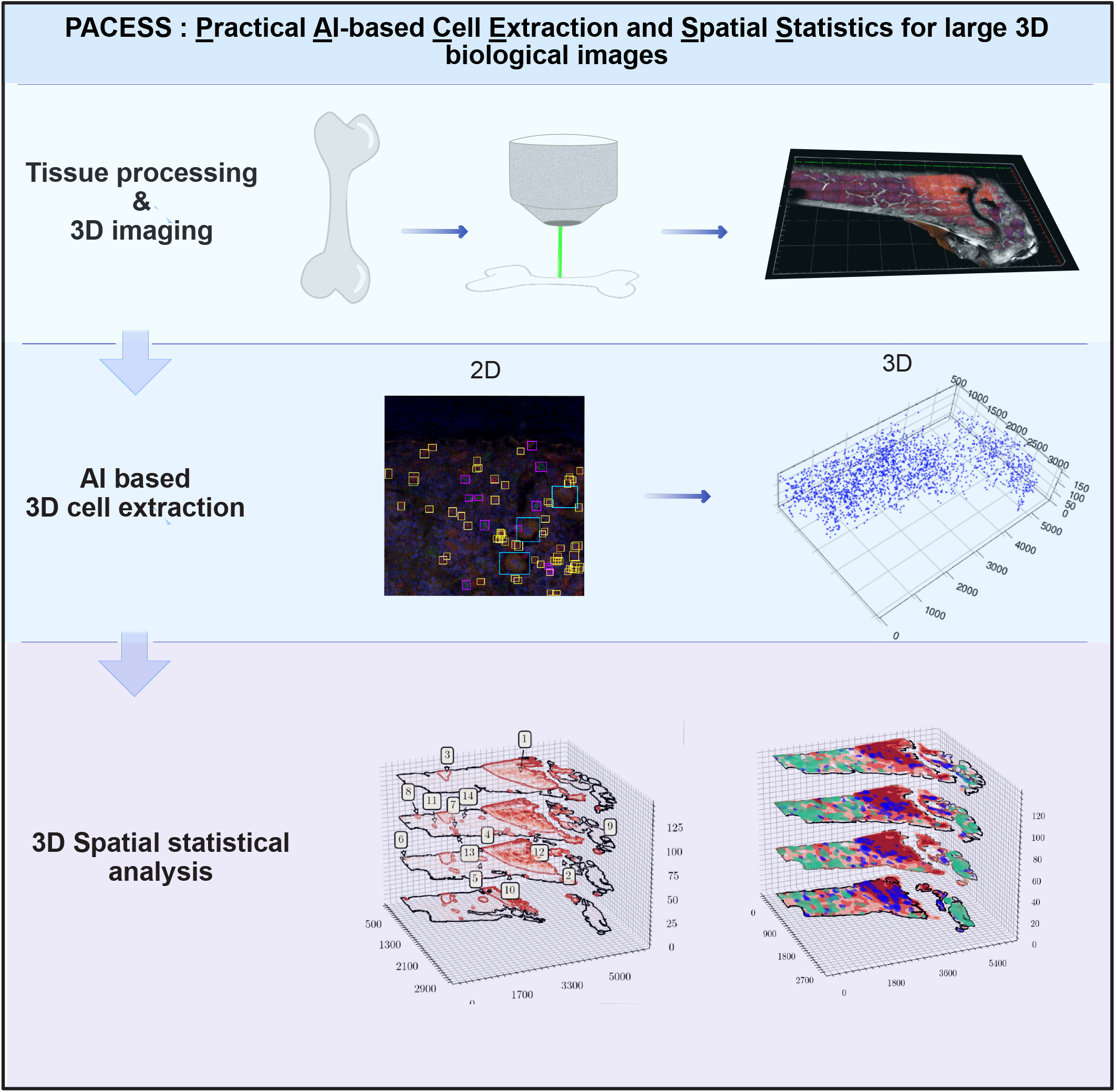

## INTRODUCTION

The cellular architecture of organs is known to be a key feature for the maintenance of their function, and changes of cellular organization are often linked to disease. This has been extensively demonstrated for tissues that have well-known structures such as the skin or the brain ^1,2^. The Bone Marrow (BM) is the organ where hematopoiesis, the process of maintaining all blood cells’ turnover through the daily generation of billions of diverse cell types in mouse, and trillions in human, takes place. The combination of flow cytometry, single cell transcriptomics and functional assays has provided a wealth of information on the different hematopoietic cell populations co-existing in the BM ^3–5^. However, still very little is understood about the spatial organization of this tissue. BM is the softest tissue in the body and has long been considered as an amorphous, even liquid, structure with a wide range of different cell types tightly packed within a confined cavity. These features make the BM a uniquely challenging tissue for histological imaging and quantification. To address this challenge, researchers in the field have made use of increasingly sophisticated BM multidimensional imaging techniques (3D or greater) focusing on visualizing hematopoietic stem and progenitor cell (HSPC) populations in large BM tissue preparations ^6–8^. While these led to the emerging concept that the BM is spatially and regionally organized to differentially support distinct HSPC populations^6,9^, the spatial organization of multiple hematopoietic cell types remains unclear. Understanding it holds clues for more effective harnessing of HSPC function, development of improved therapies for hematological disorders and overall healthier ageing. Quantification of multicellular interactions within large BM 3D image datasets has not yet been achieved for two main reasons. First, image segmentation by intensity thresholding, which is, historically, the most widely used method for identifying cells of interest, underperforms in scenarios where cells are compressed or tightly packed together, as in the BM, where boundaries between cells become more difficult to distinguish. Hence, a high number of cells are not identified and are lost. The second challenge relates to the statistical methodologies available to assess multicellular interactions. Inferences have typically been made using pairwise hypothesis testing, but this approach is limited as it does not facilitate the direct examination of the relationships between a higher number of cell types, making it challenging to formulate definitive conclusions from the underlying data.

Here we present PACESS (Practical AI-based Cell Extraction and Spatial Statistics), a pipeline for extracting and analyzing hematopoietic cells from large 3D BM images, overcoming existing limitations (graphical abstract). This pipeline makes use of convolutional neural-network based object-detection to classify and identify the locations of cells in 2D, followed by an automated method that creates 3D understanding. We introduce an augmented object detection deep neural network trained using 2D data alone, for which images can be rapidly annotated. Annotations are created for each image-layer in the 3D data and the output from multiple layers are automatically combined to identify each cell’s type and location within the 3D space. Once the spatial data are extracted, we apply a set of 3D spatial statistical analyses. The steps in this process include: generating exploratory statistics that quantitatively assess spatial heterogeneity; identifying regions of high cellular density of specific cell types; visualization of the local density of cell types; and, finally, identification and visualization of location-dependent relative abundance between cell types. The resulting analysis is presented as a holistic ‘statistical map’ of BM tissue.

We applied PACESS to study the spatial interactions of megakaryocyte and T cells in the context of leukaemia. Leukaemia progression leads to healthy hematopoietic cells being dislodged ^9,10^, and therefore represents a good model for studying changes in multiple cell populations’ spatial interactions. First, PACESS was able to efficiently extract the cellular information for the three cell populations, even in areas tightly packed with leukaemia. Then, the statistical analysis revealed that the density of AML cells at a specific BM location significantly affects T cells and megakaryocytes’ densities, albeit with distinct patterns, resulting in the generation of multiple microenvironments characterized by unique relative densities of these three cell types.

## RESULTS

### Large 3D image datasets of murine BM represent a computational challenge for analysis

By nature, the bone marrow (BM) is an organ where cells are highly packed together. This is exacerbated during acute myeloid leukaemia (AML), when malignant cells outcompete healthy hematopoiesis through mechanisms that are still not fully understood. To study the spatial relationship of different cell types within the BM, it is informative to work with large 3D tissue specimens. We generated these using agarose embedding and a vibratome to cut 200-300 μm thick sections of various bones, a combination of elements from previously described clearing protocols developed for BM or other tissues, and careful troubleshooting of tissue fixation (Figure S1).

Heme is highly abundant in the BM and is known to interfere with light penetration, generating background noise and thus reducing image quality. We first used the Reagent 1 (or ScaleCUBIC1) solution from the CUBIC method to decolorize heme efficiently^11^. To quickly clarify BM samples while maintaining their integrity over time, we selected the Ce3D tissue clearing reagent and kept the samples in it for imaging and storing^12^. Finally, we embedded samples in Histodenz reagent, resulting in the generation of transparent samples (Figure S1A-B). To confirm that our clearing protocol did not distort BM tissue, we imaged the calvarium BM cavities of a vWF megakaryocyte transgenic reporter mouse using two photon excitation intravital microscopy (without any clearing) first, and then again following explant and clarification. The mouse was injected with FITC-conjugated dextran to identify vasculature during IVM, and the calvarium sample was stained with anti-endomucin antibody to achieve the same result *ex vivo*. Juxtaposition of the tilescans obtained via IVM and following clearing (Figure S1C) revealed no changes in the overall appearance of the tissue. Moreover, quantification of the diameter of megakaryocytes demonstrated no difference in the two images (Figure S1D). Finally, to maximize the signal to noise ratio in thick BM sections we tested the effect of fixation on immunostaining clarity. While we found fixation length affected some antibodies more than others, we identified 2 hours fixation as the ideal, goldilocks timing that allowed both efficient vibratome cutting and easily interpretable immunostaining results (Figure S1E-F).

To generate our exemplar image, we injected a wild type C57Bl/6 mouse with tomato+ AML cells generated through retroviral transduction of the oncogene MLL-AF9, a well-established and widely used murine model of AML ^13,14^. We harvested bones at an intermediate disease stage, whereby flow cytometry analysis confirmed an overall infiltration of femur bone marrow of 15%. A 250 μm thick section from the contralateral femur was stained with an anti-CD8 antibody followed by species specific fluorophore-conjugated secondary antibody and DAPI nuclear counterstain to highlight cytotoxic T lymphocytes (CTLs), cleared and imaged using a combination of confocal and two photon microscopy. In the resulting 3D tilescan image, bone was visualized through second harmonic generation (SHG) signal of collagen (represented in grayscale), and AML cells, CTLs and megakaryocytes (MKs) were identifiable through their membrane expression of tomato fluorescent protein (red), anti-CD8 immunostaining (green), and characteristic morphology (large size, multilobate nucleus) and autofluorescence (pale green/red signal), respectively. Moreover, all nuclei were visible thanks to DAPI counterstain (blue) (Figure 1A).

**Figure 1:**
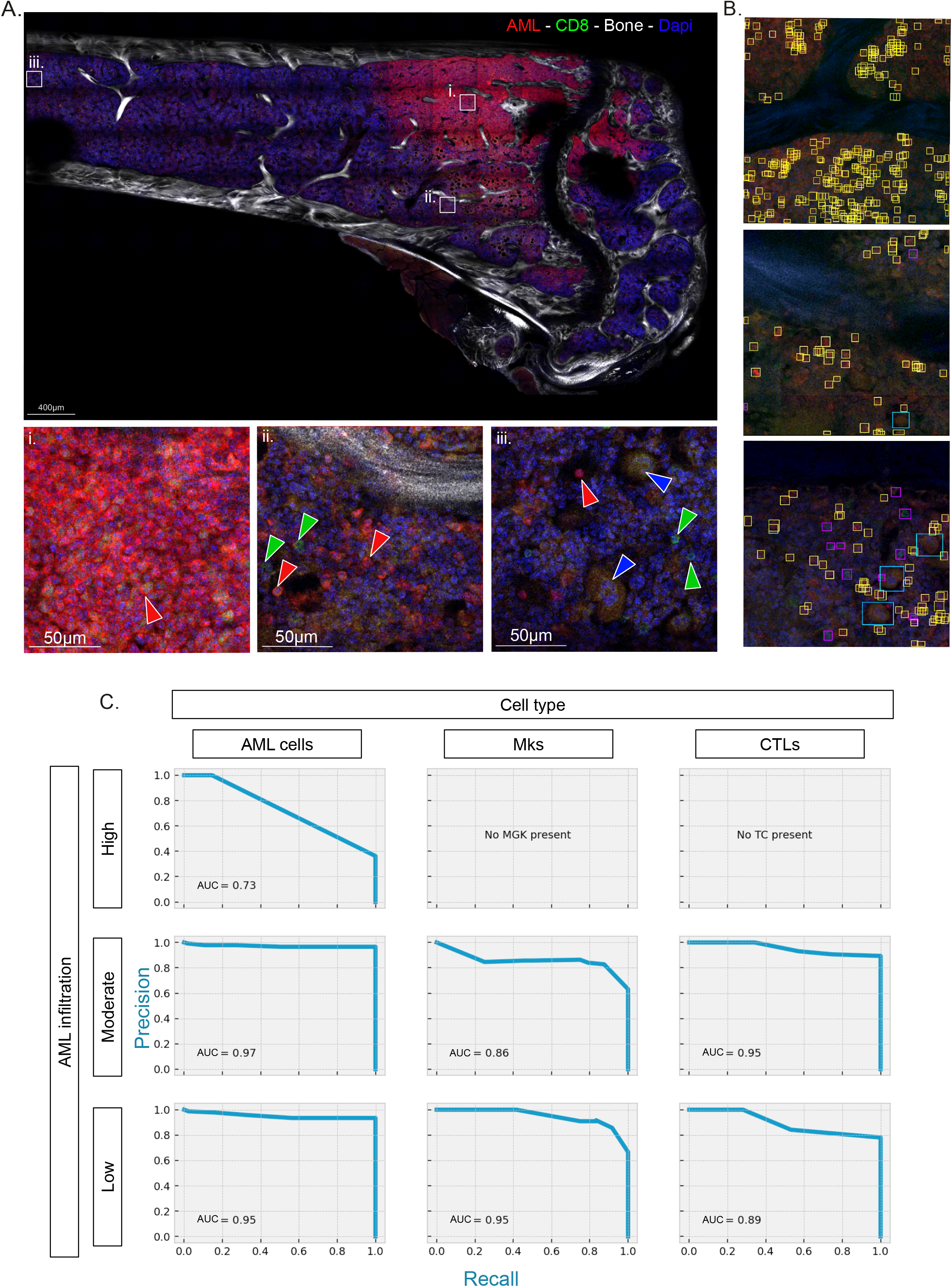
2D data extraction using deep neural network. **(A)** Tile scan of a single z position (z=27 out of 32) of a clarified 250µm thick section of tibia from AML infiltrated mouse. CTLs’ cell membranes are labelled by CD8 immunostaining (green), AML cells expressed membrane-targeted tdTomato (red) and low level of intracellular GFP (green), DAPI counterstained nuclei (blue), bone is image using SHG (grey). Mk are identified thank to their specific cellular morphology (large size, multilobate nucleus and autofluorescence). **(i-iii)** High magnification images of specific area with high, intermediate and low AML infiltration respectively. Arrows show representative examples of AML cells (red), CTLs (green) and MKs (blue/white border). **(B)** Example of AML (yellow boxes), CTLs (purple boxes) and MKs (blue boxes) predictions from the 2D object detection. Shown are example images of highly (top), intermediate (middle) and low (bottom) AML infiltrated areas. **(C)** Precision-recall curve from the 2D object detection model for each cell type in area of high (top panels), intermediate (middle panels) or low (bottom panels) AML infiltration.

An initial inspection of the image confirmed the uneven distribution of AML cells, whereby the BM region near the growth plate was very heavily infiltrated by AML cells, tightly packed together (Figure 1A i), whereas the remaining tissue displayed varying levels of local AML infiltration (Figure 1A ii and iii). However, it was evident that it would be a challenge to quantify local differences in the density of T cells and megakaryocytes, as well as subtle differences in AML cell density, and, even more, to identify whether the density of each cell type may be a function of the density of one or both the other cell types. Such work is important to understand the cellular dynamics taking place in BM tissue as AML cells invade it, and relies on the precise identification and annotation of all the cell types of interest.

### Data extraction using deep convolutional neural network

The image we generated is 15 GBs large in size and is the result of the juxtaposition of 15,776 partially overlapping 2D images acquired as adjacent z stacks, precisely 32 2D tiles of 493 512×512 pixel tiles, with a z dimension step size of 5 μm. Traditional image analysis software would open the entire image in order to process it and are not equipped to process 3D images of this size. We therefore selected a convoluted neural network model that could be trained using small 2D image samples randomly selected from the overall tilescan, would operate on image portions and generate output data of much smaller size and therefore easy to handle computationally^15,16^. We manually annotated a total of 18,240 cell examples, of which 14,655 were AML cells, 3,056 CTLs and 529 MKs. 70% of these annotations were used for training, 20% for validation and 10% for testing. After 200 iterations (epochs), we identified the optimally trained network as the one generated after 60 epochs. This fully trained network annotated 506,611 cells across all 2D slices using bounding boxes that could be easily inspected by eye (Figure 1B). Next, to measure the model’s accuracy, we manually classified all test images according to whether they represented areas of high, moderate or low AML infiltration (e.g. Fig. 1Ai, ii and iii, respectively) and we compared our manual annotation with the results of the model to generate precision/recall curves (Figure 1C). Areas where AML cells were tightly packed together were the ones where the model performed worst in identifying AML cells themselves (Figure 1C, top left panel), even though it still returned high numbers of AML cells (e.g. Figure 1B, top panel). In these same areas CTLs and MKs were identified efficiently by the model (Figure 1B), however there were too few of them to generate meaningful precision/recall curves. In areas of intermediate and low AML infiltration the neural network achieved a mean AUC 0.8942 (Figure 1C), with AML cells, CTLs and MKs all accurately identified consistently (see respective AUCs in Figure 1C panels). In conclusion, the model was able to identify cells in 2D images based on a combination of fluorescent signals and morphology features.

To generate maps of the 3D locations of cells across the entire dataset, neighboring boxes from across different image layers were linked together using a 3D aggregation algorithm we designed. This was an important simplification of the workflow because these 3D maps could be generated without the need for 3D labelled images, however the resulting 3D boxes could be viewed in samples of the original image (Figure 2A-C). To test the accuracy of this step, we randomly selected 3 separate 210×210×50 μm 3D regions with high, medium and low AML infiltration within the overall image, manually annotated AML cells, CTLs and MKs in 3D, and compared manual and predicted annotation (Figure 2D). Importantly, we didn’t have a large deviation from the diagonal when we plotted the results of the two methods, with only a slight underestimate of cell counts by the model compared to manual classification found at higher cell densities. Using the automated approach, we identified 197,726 AML cells, 8,225 CTLs and 2,160 MKs, which would be infeasible manually. In addition to their number, their specific localization was extracted by this analysis (Figure 2E-G).

**Figure 2:**
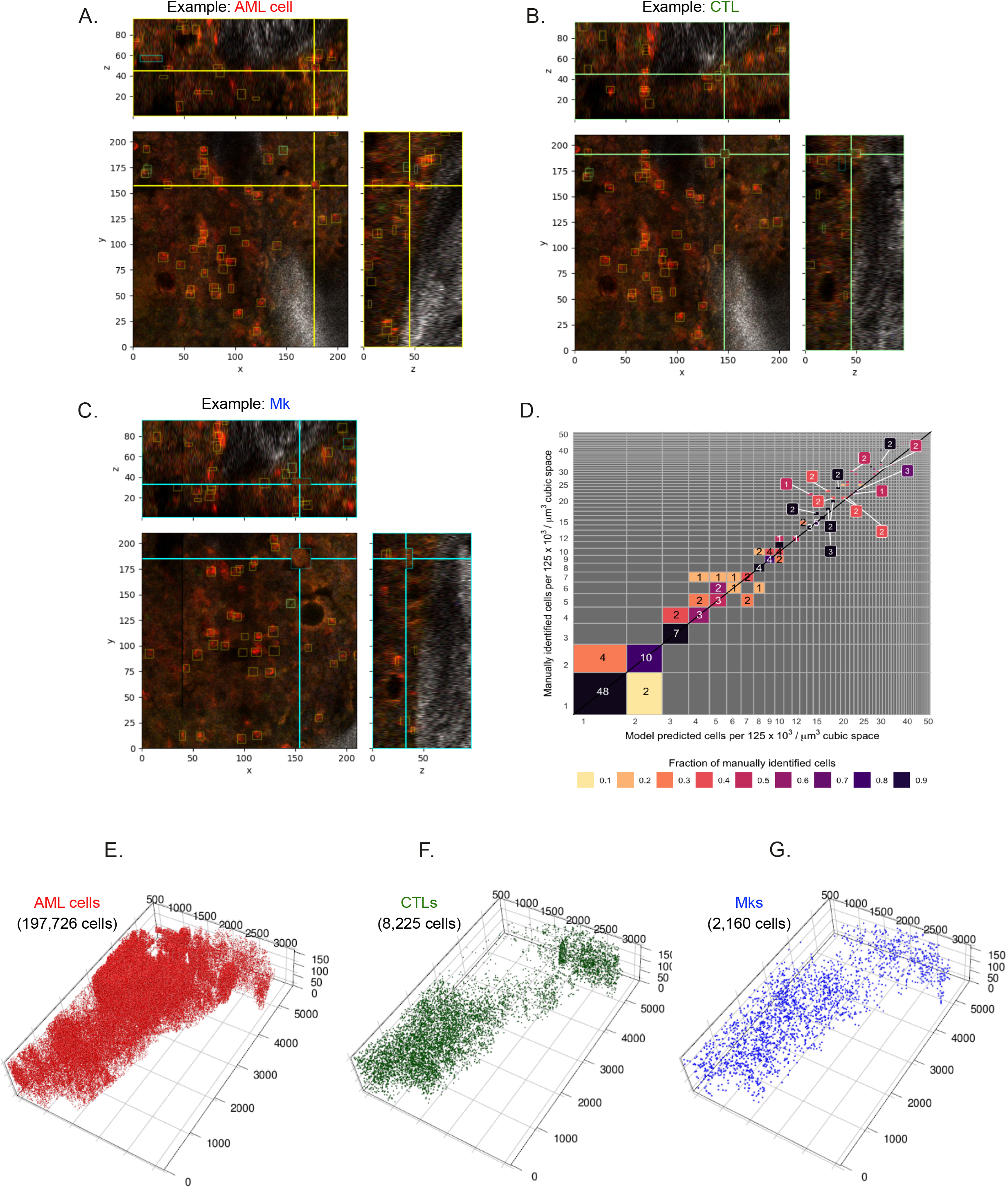
3D reconstruction of the bone marrow cellular organisation. **(A-C)** Example images of 3D bounding boxes for AML cells (**A** – yellow boxes), CTLs (**B** – green boxes) and Mks (**C** – blue boxes). AML cells expressed membrane-targeted tdTomato (red) and low level of intracellular GFP (green), CTLs’ cell membranes are labelled by CD8 immunostaining (green) and Mks are identified thank to their specific cellular morphology (large size, multilobate nucleus and autofluorescence). Scales are in µm. **(D)** Comparison of manually identified vs model predicted cells identification per 125 ×10^3^/µm^3^ cubic space. 3D reconstruction of AML cells **(E)**, CTLs **(F)** and Mks **(G)** localisation in the bone marrow. The total number of identified cells for each population is indicated in black.

Visualization of all identified cells highlighted how 2D slices of BM tissue can lead to an underestimate of the prevalence of AML cells even in areas that appear to be little infiltrated (Figure 2E). Consistent with previous reports of healthy hematopoietic cells being excluded from highly infiltrated areas of BM tissue^14^, the area near the growth plate appeared devoid of CTLs (Figure 2F), however MKs seemed to present a different distribution pattern from that of CTLs (Figure 2G), highlighting the need for a dedicated pipeline of spatial analysis that would allow better understand the spatial organization of each cell type and the spatial relationships between cell types.

### Spatial heterogeneity and automated identification of areas of high cellular density

To visualize and analyze the spatial distribution and relationships between AML cells, CTLs and MKs, space was discretized into non-overlapping, adjacent cubes that cover the entire 3D area, and are the units analyzed using spatial statistics. The latter were first used to quantify heterogeneities in the distribution of cell types and, next, to explore cells’ interdependencies. To assess spatial heterogeneity we used two steps: first, we used Moran’s I (also known as spatial autocorrelation function) ^17^, to assess the similarity of adjacent cubes; next, we linked similar cubes to outline areas of high cellular density using DBSCAN^18^ (Figure 3A).

**Figure 3:**
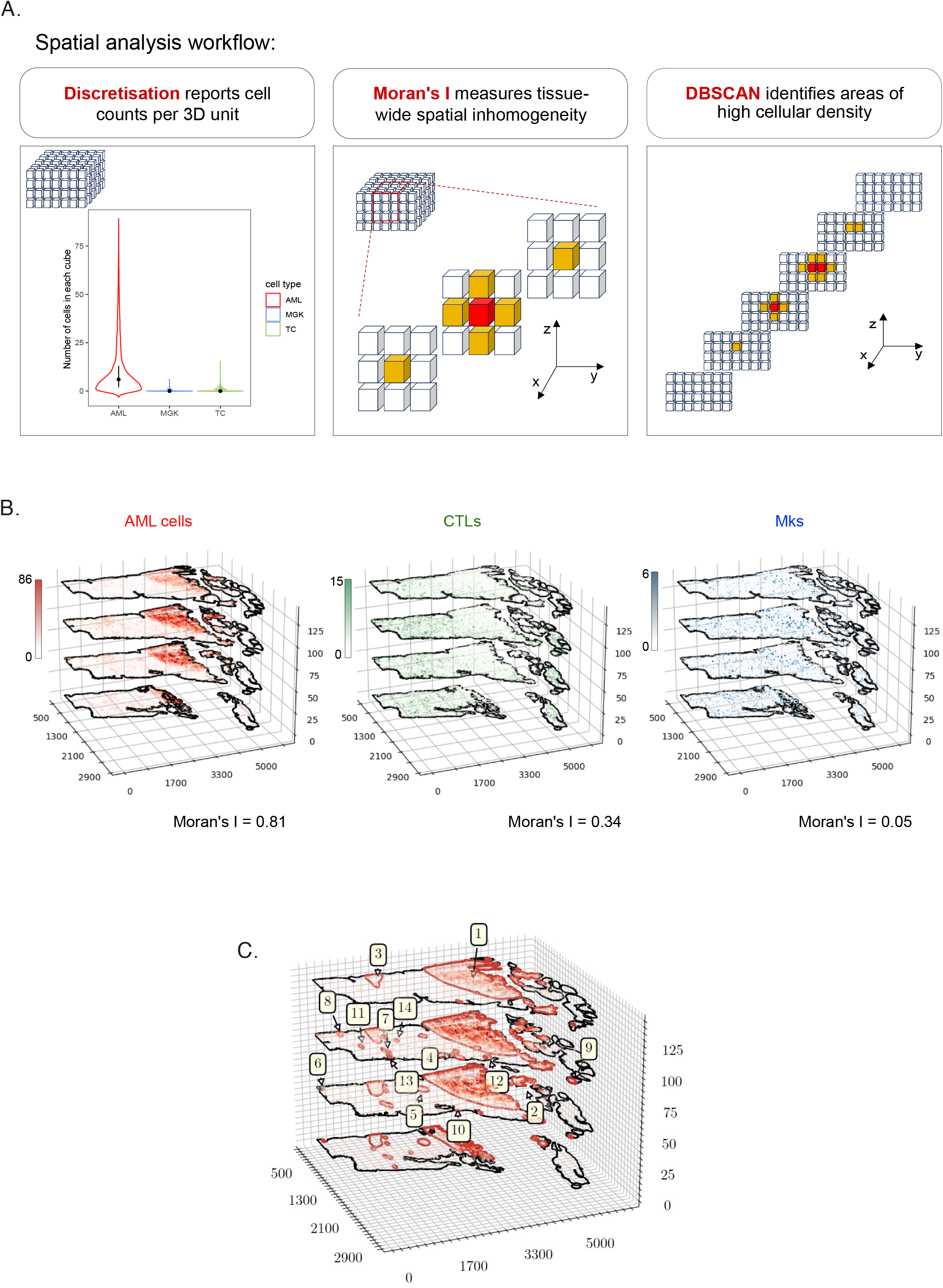
to be completed once final figure agreed. **(A)** Summary of the spatial analysis framework. Discretisation of space allows recording cell densities. Moran’s I measures the spatial autocorrelation of each cell type. In the data, the neighbours of a cube (red) are those immediately adjacent (yellow). Moran’s I uses that local geographic information to calculate a global measure of spatial autocorrelation. DBSCAN is used to identify areas of high cellular density. The centre cube (red) and its neighbours (yellow) are considered a local high-density region if, taken together, they contain more cells than 7 (corresponding to the number of cubes) times the third percentile per cube for that population. Adjacent local high-density regions are grouped to form a contiguous cluster. **(B)** Quantitative 3D visualization displays the number of AML cells, CTLs and MKs at each location, with colour scales indicating abundance. Moran’s I values are reported for each cell type. **(C)** 3D projection visualization shows AML clusters identified by DBSCAN. The clusters are numbered in decreasing order of total AML cell number contained in them.

The first step was to identify the ideal cube size to work with. For meaningful visualization, the discretization needs to be such that some aggregation of cell counts occurs, but a few cells of the same type are found in each cube. A cube size of 45 μm^3^ was selected to be sufficiently fine that geographic resolution was retained, but sufficiently coarse that the resulting data could still be computationally assessed without undue burden. The number of cells of each type in each cube was recorded (Figure 3A, left panel), and with that discretization we could visualize the density of AML cells, CTLs and MKs as a function of their position (Figure 3B). This information was the basis to quantify spatial heterogeneity in the distribution of each cell type and of combinations of cell types.

To quantitatively assess the homogeneity of each cell type’s spatial distribution, we calculated Moran’s I. To calculate Moran’s I, one must consider the similarity between each spatial unit (one of the cubes) and its neighbors. Neighbors can be defined in several ways and Figure 3A, middle panel, illustrates our definition of neighborhood, which is well-suited for the following steps of our spatial statistics pipeline. Briefly, when cubes had one edge in common, they were considered to be neighbors, resulting in each cube being the central one of a neighborhood of seven neighboring cubes where six have a face in common with the central one (Figure 3A, middle panel). Moran’s I is a measurement of the overall homogeneity of cell numbers in cubes across the tissue with reference to their neighbors. If Moran’s I is positive, cells tend to be aggregated in common areas. If Moran’s I is close to zero, cells are distributed randomly in space. When its value is less than zero, cells are more homogeneously dispersed than one would expect from a random process. Moran’s I for the AML cells was 0.81, quantitatively substantiating the observation that AML cells were concentrated in relatively few areas. Moran’s I for CTLs was 0.34, suggesting some positive spatial clustering but less than found for AML cells. Moran’s I of 0.05 for MGKs indicated these cells had a weak geographical dependency, resembling that of random locations (Figure 3B).

While Moran’s I can indicate that cells of a given type are overall co-located or scattered throughout the tissue, a distinct methodology is needed to identify tissue regions where those cells are at high density. To do this, we used the Density-Based Spatial Clustering of Applications with Noise (DBSCAN) algorithm^18^. If the average number of cells in the cubes of a 7 cubes neighborhood was greater than the third quartile value for that population in all cubes across the sample, whereby only a quarter of cubes have more cells than this value, the neighborhood was considered dense. This calculation would be repeated by shifting the neighborhood center from one cube to one adjacent one, until all possible neighborhoods would be considered. As a result, dense neighborhoods were agglomerated to highlight contiguous areas spanning cell clusters (Figure 3A, right panel).

For the AML data, DBSCAN identified 18 distinct high-density areas, or clusters. The largest 14 are marked in decreasing order from largest to smallest (Figure 3C). The largest cluster accounted for 61.8% of all AML cells in the image, and the following two largest for 5.9%, and 2.0% of all AML cells. The two largest clusters of AML cells, 1 and 2, were proximal to the growth plate, on either of its two sides. Cluster 3 was more distal, and the overall the distal region of BM contained much smaller clusters. Interestingly, larger clusters were located adjacent to the endosteum, while centrally located clusters tended to be smaller. For CTLs and MKs, respectively, approximately 70% and 90% of cubes recorded a zero cell-count (Figure 3A, left panel), and, consequently, DBSCAN returns no clustered areas. This was consistent with these cells being less abundant than AML cells, generally not clustered and returning Moran’s I values that were positive but close to zero.

### Spatial dependencies between cell types

The key to understanding the principles regulating multicellular tissues is based on understanding how diverse cell types interact with each other to enable tissue function. It is therefore important to explore whether local heterogeneities in the distribution of specific cell types depend on those of other cell types, and to identify and map the resulting different cellular neighborhoods. To do this, we first asked whether, besides low Moran’s I and lack of clustering, CTLs and MKs localization could be described as a function of AML cell presence and/or density. To investigate this, we used permutation tests and logistic regression (Figure 4A) ^19^.

**Figure 4.**
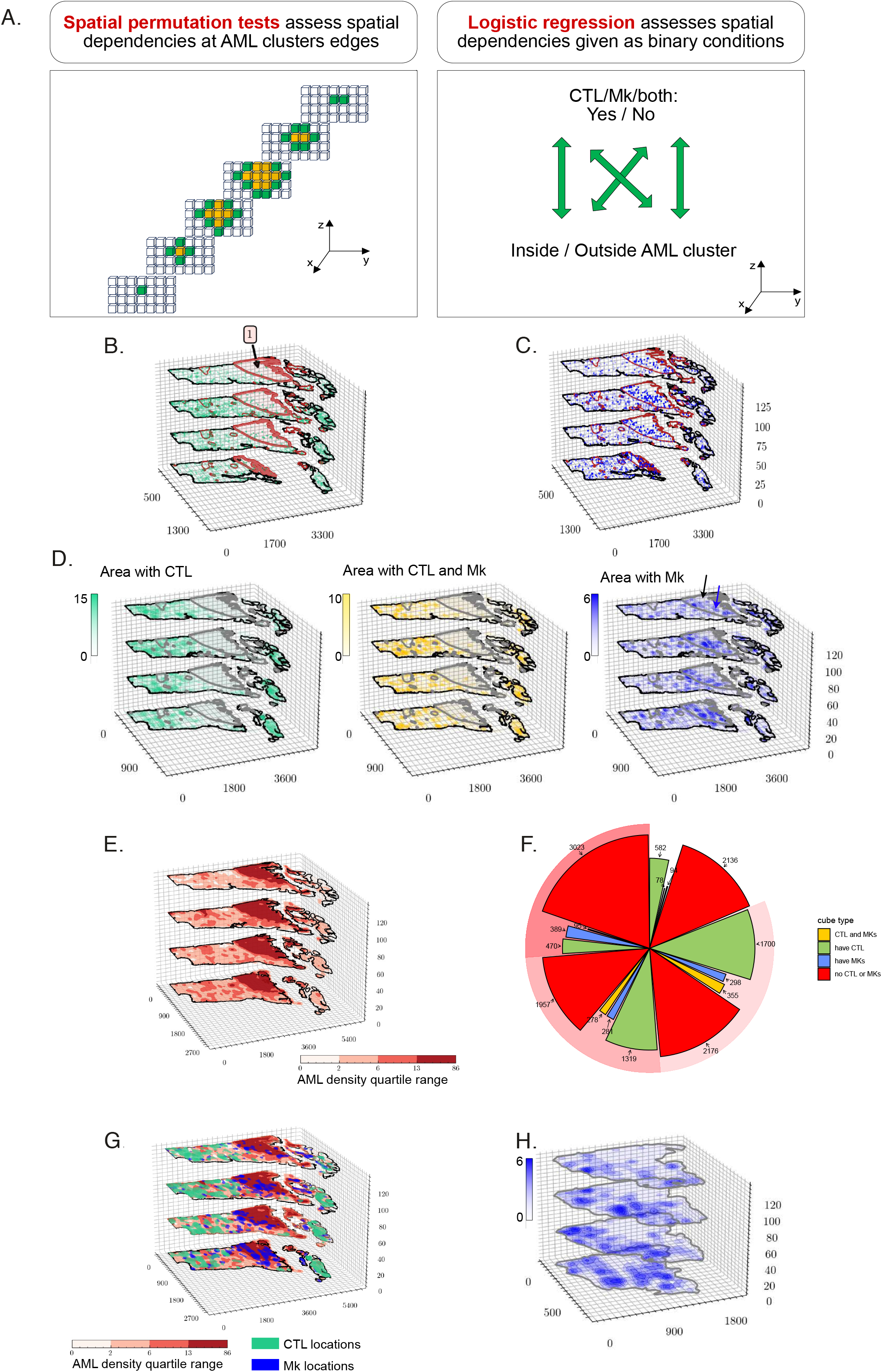
Spatial relationships between cell types. **(A)** Left: for each cluster of cubes (yellow), the cubes surrounding them are identified (green). Spatial permutation tests are then employed to challenge the hypothesis that the mean density of cells of a second type is independent of whether those cells found within the cluster or in the area surrounding it. Right: Logistic regression was performed to assess whether the presence or absence of other cell types was independent of dense areas. Statistical significance of each binary variable was assessed using t-tests. **(B)** 3D projection visualization shows hypothesis test results for CTLs. The null hypothesis for CTLs is rejected for the largest cluster with Holm-Bonferroni corrected p=0.00007. **(C)** 3D projection visualization shows hypothesis test results for MKs, where all null hypothesis are not rejected with p-values >0.05 after Holm-Bonferroni correction. **(D)** Results of logistic regression modelling. Likelihood of finding CTLs (left), CTLs and MKs (middle), and MKs (right) within dense AML areas. CTLs p value is <2e-16. CTLs and MKs p value is <2e-16. MKs p value is 0.464. Shades of colours highlight density of each cell type. Arrows in F highlight areas within AML dense area 1 that are MK rich (blue arrow) or devoid (grey arrow). **(E)** Cubes were labelled based on quartile distribution of AML density (left). **(F)** Cubes were further labelled within each AML density quartile based on binary information about containing CTLs, MKs or both. **(G)** Information on the presence of CTLs and MKs was overlaid to the AML quartile information. **(H)** Higher magnification rendering of dense AML area 1, highlighting MK rich and devoid areas, with shades of colours indicative of MK density.

For CTLs, visual inspection of the image indicated that their density was uneven throughout the tissue, and there were very few CTLs in the area containing the largest AML cluster, cluster n.1 (Figure 2E-F and 3B-C). To statistically assess if the distribution of CTL counts was influenced by any areas of high AML density, permutation tests were used to challenge the null hypothesis that the mean number of CTLs within each cube is independent of whether the cube is within an AML cluster or adjacent to it (Figure 4A, left panel). The null hypothesis was rejected only for the AML cluster 1 area, highlighted in Figure 4B. The remaining 17 smaller AML clusters all failed to reject the null hypothesis. This analysis indicated that only the largest AML cells high density area influenced the mean number of CTLs per space unit. In other words, CTLs were generally missing where the largest AML cluster was located, but their density was not affected in the smaller areas with dense AML cells.

Spatial permutation tests were also performed to test the null hypothesis that the mean number of MKs in each cube is independent of whether the cube is within an AML cells high density area or adjacent to it. The results showed no evidence that MKs had a different distribution within versus around any AML dense area (Figure 4C). However, visual inspection of the image highlighted how the largest dense AML area could be further split into two sub-regions based on higher and lower density of MKs, indicating that further spatial statistical tests and visualization tools were needed.

To measure the likelihood of finding CTLs, MKs, or both cell types in dense AML areas compared to non-dense AML areas, we used logistic regression, a statistical model that reports on the likelihood of an event to occur or not given a certain binary condition, in our case the presence of CTLs, MKs or both inside vs. outside dense AML areas (Figure 4A, right panel). The likelihood of finding cubes with CTLs or both CTLs and MKs inside dense AML areas was 80% lower than that of finding such types of cubes outside of dense AML areas (both p values <2e-16). The likelihood of finding cubes with MKs but not T cells inside dense AML areas was lower, but not significantly, than outside of them (Figure 4D). For MKs, this latter result was driven by heterogeneity within dense AML area 1, where a sub-area was present that was devoid of MK cells, while a large proportion of this area still contained MKs, and MKs there appeared to be denser than in the rest of the image (Figure 4D, right panel, black and blue arrows, respectively).

To highlight how AML density alone would not be sufficient to determine whether MK density would be affected, we developed a visualization tool whereby we binned all cubes based on their quartile AML density values (Figure 4E. Next, we further classified each cube according to whether it contained T cells and/or MK cells. This resulted in 16 possible cellular composition types based on AML density and presence/absence of CTLs and MKs. While all 16 possible neighborhoods existed, some were more abundant than others, with areas containing both CTLs and MKs being rarest, independently of AML density quartile (Figure 4F). Spatial visualization of the neighborhoods highlighted how outside of dense AML area 1 various types of neighborhoods could be found adjacent to each other, without a specific pattern (Figure 4G). The dense AML area 1 was split in two regions based on MK presence. Even though the MK-rich area was closer to the distal part of the bone, and the area closest to the growth plate was generally devoid of MKs, the MK-rich and -devoid areas were not simply adjacent, because MK-containing areas were most often surrounded by MK-depleted areas (Figure 4H). Both inside and outside AML dense area 1 MK-rich and CTL-rich areas were irregularly shaped, with more jagged contours than AML dense areas, indicating that while regional variations in tissue composition could be identified, these regions had irregular and intertwined shapes.

In conclusion, our pipeline adapts spatial statistics to a 3D framework to enable the quantitative evaluation of clustering, the automatic identification of regions of high density of specific cell types, and the statistical assessment of dependencies between cell types to map unique cellular neighborhoods characterized by unique combinations of densities of multiple cell types. It allowed to identify that only very large BM areas densely infiltrated by AML cells are devoid of CTLs, and that these areas can be further subdivided according to the density of MKs. Moreover, the transition between area types tends to be gradual, with areas exhibiting rough, jagged borders.

## Discussion

In this study we have described a novel, efficient and effective pipeline, PACESS, for extracting meaning and drawing inferences from large 3D BM tissue images. These images are relatively affordable to generate, but very challenging to analyze efficiently because of their size and complexity. It has been especially challenging to understand and quantitatively evaluate the relative spatial organization of cell types. This type of analyses, however, holds the promise to allow a much deeper understanding of the cellular principles regulating hematopoiesis, with important implications for the development of better therapeutic and preventative strategies for hematological disorders.

In terms of tissue preparation for image acquisition, the choice of clearing and mounting solutions was critical to maximize tissue integrity including over time. While other published solutions led to tissue darkening and loss of fluorescent signal within a few days^7^, our samples could be re-imaged multiple times including weeks apart. Fixation was the other critical parameter. Here a short, two hours only, fixation step proved to be the optimal solution to facilitate tissue cutting while achieving the highest signal to noise ratio.

The image presented here is a 3D tilescan composite of 15,776 images, resulting in a >4×10^9^ pixels and 15 Gb large file, and could not be opened using gold standard open access image analysis software such as ImageJ. Our image analysis workflow builds on established techniques from machine learning and geospatial statistics and has several advantages over current approaches. From the perspective of data-extraction, the workflow is reliable and scalable, and importantly solves the challenge of efficiently labelling cell types in 3D by only necessitating labelling in 2D followed by automated methods to generate 3D understanding. The spatial statistics workflow allows identifying areas where any cell type of interest is denser than average and highlights tissue regions with unique cellular composition.

While AI-based extraction methods have been applied in other contexts, murine bone marrow tissue provides distinct challenges because hematopoietic cell types are tightly packed together in an amorphous structure where no clear patterns have been identified. For 2D extraction, the YOLO neural network model was ideal because it proved to be highly effective while minimizing time (approximately 40 minutes for images of similar size to the one presented here) and computational power required. As images are provided to the model in their raw form, avoiding extensive image processing allows further time savings. While we maximized the efficiency of data-extraction by running the model on a high-performance computing cluster, we were able to run the model on cloud-based computing resources, including free ones, such as Google co-lab, without the need for additional computational resources. These characteristics provide the opportunity to analyze both larger samples, images acquired at higher resolution, and images containing a higher number of cell types.

A known drawback of AI-based approaches is the need to generate manually annotated training data. Important for the classification of hematopoietic cells, the model was robust in identifying cells with irregular/variable morphology such as megakaryocytes, and was reliable in cases where a cell type was very abundant, resulting in highly compacted, adjacent cells, such as AML cells^15,20,21^. While we demonstrate the practical application of the model to three cell types, 2D neural networks such as YOLO are scalable, and can be further trained, shared and updated to include further cell types^15^, while our method to create 3D meaning is efficient regardless of the number of cell types. The effort required to train and optimize a neural network is however outweighed by the high returns achieved when applying the network to several images. Moreover, pre-trained models can be updated with less extensive datasets through a process called transfer learning^20,21^. Such ‘pre-trained’ models become akin to the availability of deposited gene signatures greatly increasing the accessibility and productivity of bioinformatics analyses of genomics data. The dataset presented here is already a pre-trained model and can constitute the starting point for identifying other cell types in the BM but also in virtually any tissue. Of note, as 3D image analysis software is still unwieldy, grouping of the neural network 2D results into 3D predictions is a simple solution to the widespread challenge of annotating and analyzing 3D datasets.

Another important solution was utilized to build our pipeline of spatial statistics analysis. In previous studies, researchers have challenged the null hypothesis that the distribution of each individual cell population is uniform-at-random throughout the space through the use of a simulation method called random dots. In a spatial analysis context, however, the null hypothesis must be conditioned to take values only within the observable, imaged, space. Although the random dots method is intuitive, there is no guarantee that the null hypothesis is physiologically viable as we typically cannot observe all features within the tissue with current technology, which introduces a risk of bias in the findings by increasing the likelihood of rejection. We circumvent this shortcoming in our spatial statistical pipeline, avoiding the use of simulated data, and instead address the question of how cells within the bone marrow are related by other approaches, including direct comparisons between multiple different cell populations.

Our spatial statistics approach provides a reproducible method for quantifying properties of individual cell types, as well as interactions between them. Simple measures evaluate the extent of spatial clustering of individual cell types, while geographically aware clustering methods then identify regions of high density. Permutation tests provide statistical measures for assessing the relatedness of cell densities in reasons of interest, and logistic regression quantifies the extent to which the relative abundance of cell types are contingent on others. Importantly, while we demonstrate our spatial analysis workflow with three cell types, this could be scalable to include any number of cell types.

While interest in leukaemia and T cells interactions has been growing, stemming from immunotherapy developments, megakaryocytes are not normally studied together with the other two cell types, and our image analysis highlighted some unexpected spatial relationships between the three. Consistent with previous reports of AML cells growing in localized patches within the bone marrow tissue^10,14^, several clusters of AML cells were identified. All larger clusters were adjacent to endosteum, again consistent with previous reports that the AML model we analyzed tends to grow from endosteal regions^14^. The fact that no cell type returned a negative Moran’s I value is consistent with the current working hypothesis that hematopoietic cells are randomly distributed across BM space, with no ordered topology. Interestingly, while several reports describe severe loss of all hematopoietic cell types as a consequence of AML growth, our analysis highlighted that heavily infiltrated AML areas may not be homogeneous, as the BM area containing the largest AML cells cluster we observed could be split into two subregions, one devoid of both CTLs and MKs, the other devoid of CTLs only. This observation opens the questions whether there may be major differences in the rate of loss of different healthy hematopoietic cell types in the BM as AML grows, and whether areas more greatly devoid of healthy cells represent areas where AML cells have been present for longer.

In summary, the ultimate goal of quantitative 3D imaging is to informatively summarize, in a numerical form, the vast amount of spatial and cellular information present within an image. This is particularly useful for settings where the principles regulating cell distribution are not clear, such as in BM tissue. The work presented here provides a framework for the data extraction and analysis of complex 3D BM tissue images. Importantly, this pipeline enables analysis of images which would have proved too challenging to standard methodologies because of their size or complexity. The PACESS analytical workflow combines cell labelling and spatial statistical analysis to aid in the interpretation of spatial cellular data, which can provide a comprehensive insight of the relationships which exist within cell types in the BM. It promises to uncover the principles regulating the cellular organization of the cells responsible for the lifelong production of blood cells and their deregulation when hematological disease occurs.

## MATERIALS AND METHODS

### EXPERIMENTAL MODEL

#### Mice

All animal work was in accordance with the animal ethics committee (AWERB) at Imperial College London, UK and UK Home Office regulation (ASPA, 1986). All mice were bred and housed at Imperial College London or Sir Francis Crick Institute. C57BL/6 WT mice were purchased from Charles River (United Kingdom). vWF tomato megakaryocyte reporter mice were a kind gift of Claus Nerlov and Sten Eirik Jacobsen (Oxford university). Male and female mice > 6 weeks old were used. Animals were housed in Tecniplast mouse greenline cages with appropriate bedding and enrichment. The temperature, humidity and light cycles were kept within the UK Home Office code of practice, with standard diet and water *ad libitum*, the temperature between 20 and 24 °C, the room humidity at 45–65% and a 12-hours light /12-hours dark cycle with a 30-min dawn and dusk period to provide a gradual change.

#### AML experimental model

Murine AML cells were generated as described^14^. Briefly, granulocyte/monocyte progenitors (GMPs) were purified from mT/mG mice, transduced with pMSCV-MLL-AF9-GFP retroviruses as described^13^ and transplanted into sub lethally irradiated mice (two doses of 3.3 Gy, at least 3 hours apart). At 8+ weeks post-transplantation, recipient mice developed leukemia characterized by multi-organ infiltration. tdTomato positive cells were harvested from BM and spleen and cells from each primary recipient were labelled as a separate batch and cryopreserved. Primary cells from were thawed, suspended in PBS and 100,000 viable cells were injected i.v. into secondary, non-conditioned recipient mice.

#### Intravital microscopy

Intravital microscopy was performed using a Zeiss LSM 980 upright confocal microscope equipped with 5 Argon lasers (405, 488, 561,594 and 639nm), and an Insight (Newport Spectraphysics) 2-photon laser with two excitation lines of which one is fixed and one tunable (1045nm and 680-1300nm respectively), 6 non-descanned external detectors including 2 nose-piece detectors (GaASP). Images were acquired using a 20x, 1.0N.A., water immersion lens with 1.4mm working distance. Live imaging of the calvarium bone marrow was performed as previously described^22,23^. Dextran FITC and tdTomato were excited using the 488nm and 561nm laser, respectively and detected with internal detectors.

#### Immunofluorescence staining on thick bone sections

Tibias, femurs and calvarium were harvested and fixed for different period of time in 4% formaldehyde at 4°C. Bones were decalcified for 10 to 15 days in 10 % EDTA and tibias and femur were embedded in 4 % low EEO agarose (Sigma, A0169) and cut using a Leica T1000 Vibratome at depths of 250μm. All the following steps were performed under agitation at room temperature. Secctions or whole calvarium were incubated in 20% CUBIC-1 reagent (urea (25 wt% final concentration), Quadrol (25 wt% final concentration), Triton X-100 (15 wt% final concentration) in dH2O) ^11^ for 24 hours, rinse in TBS, unspecific antigen binding site were block using blocking buffer solution (TBS 0.1% Triton, 10% DMSO, 5% normal donkey serum) over nigh. The bones were then incubated with primary antibodies (CD8 clone 2.43 Invitrogen), diluted in blocking buffer for 48 hours. After several consecutive incubations in washing buffer (TBS 0.1% Triton), bones were incubated with secondary antibodies for 48 hours, followed by washes. Nuclei were counterstained using DAPI incubation over-night. Finally, the bones were mounted in the Ce3D clearing solution^12^ (Methlyacetamide 40% (sigma) diluted in TBS, 1.455g histodenz (sigma) per 1ml 40% Methlyacetamide, 4% DABCO (sigma)) using silicon isolator (ThermoFischer P18175) on Superfrost Plus™ slides and imaged at least 24 hours after, once the tissue is cleared.

#### Image acquisition

Image acquisition was performed using a Zeiss LSM 980 upright confocal microscope equipped with 5 Argon lasers (405, 488, 561,594 and 639nm), and an Insight (Newport Spectraphysics) 2-photon laser with two excitation lines of which one is fixed and one tunable (1045nm and 680-1300nm respectively), 6 non-descanned external detectors including 2 nose-piece detectors (GaASP). Images were acquired using a 20x, 1.0N.A., water immersion lens with 1.4mm working distance.

#### Object detection and clustering

The Imperial College High Performance Computing (HPC) cluster we used contains 8 CPUs and one P1000 GPU with 96 Gb of RAM and was accessed using a conventional laptop with 16GB RAM, Intel® i9-9880H CPU with a Quadro T2000 Mobile GPU.

A YOLO-V5X model (*https://github.com/ultralytics/yolov5*) was used as the backbone of the 2D object-detection neural network. More details on this model, and on convolutional neural networks more generally, can be found in the literature^15,24–26^. In this section we provide only a brief overview of the model with a focus on the manner in which the algorithm identifies objects, and how it was adapted to generate 3D estimates. Image transformations, such rotation or artificial introduction of additional noise, are only used in the training and validation stages (old ref 27) to improve the neural network’s performance when applied to unaltered experimental sample images (old ref 26).

The YOLO algorithm works by placing a multitude of boxes within the space of a 2D image and then filtering these boxes based on probability estimates from the model. It is a fully connected neural network which divides a 2D image into S × S grid of cells into which *B* bounding boxes are detected. The model identifies a set of box sizes for each class *a priori* using a k-means clustering algorithm run on box sizes observed within the training data. Each bounding box is defined by 5 parameters: the x, y central position, width (*w*), height (*h*) and a confidence score, *C*. This last value, *C*, is the confidence estimate over the presence, or absence, of an object being within the grid cell. This makes use of the intersection-over-union (IOU) between a predicted bounding box and a ground truth (manually annotated) bounding box. The greater the overlap between the two, the higher the IOU, and greater the confidence in the box. If any object is absent from the grid cell, the probability of the object (Pr(Object)) is set to 0. Otherwise it is 1. For the *i*th bounding box in *j*th grid cell, the confidence score, C_ij_ is thus calculated as^26^:

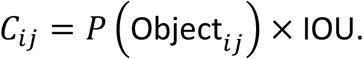

In addition to these five parameters a set of conditional class probabilities is calculated. Given *K* possible classes, this is the probability of the object belonging to any specific *k* th class: Pr(Class_k_|Object). A class-specific confidence score (CS_kij_) is then calculated as a product of C_ij_ and the conditional class probability^26^:

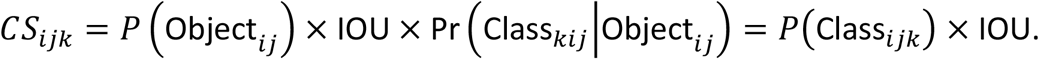

The class-specific confidence scores and IOU results are used to *select* bounding boxes through *non-maximum suppression* (NMS)^16,26^. YOLOv5 makes use of *soft-*NMS which is better adapted to overlapping objects^16^.

To aggregate the final set of 2D bounding boxes into 3D bounding ‘cubes’ we ordered the bounding boxes for each class by maximum diameter and mean fluorescence intensity (mFI). For each box within the set of boxes (*B*) within the *k*th class, starting from the largest and brightest boxes, a central x, y, z location is calculated, which we call *q*. We can also determine a maximum diameter for this box, *d*. From this point *q* the surrounding cluster of boxes in the *z* dimension which have a distance from *q* which is < d/2. We call this set of clustered bounding boxes B_c_. Within this context B_c_ ⊂ B but all the 2D bounding boxes within B_c_ are assumed to belong to a single cell (cube) surrounding an individual cell. Once identified, B_c_ is removed from the *B*. The process is repeated until every 2D box is allocated to a 3D cube.

To apply this model, an x × y × z × c dimensional image, where c = 3 for RGB, was divided into a set of 416 × 416 × 1 × 3 (RGB) tiles. Test/validate/train subsets were selected from random sampling of this tile set. Manual annotation was performed to identify cells of interest within these selected images with a minimum of 500 cells annotated within each cell class. Once trained, the final object detection was performed on the full-set of tiles.

#### Spatial analysis and modelling

Cells outside the bone marrow were excluded from the object detection dataset. Then, the bone marrow was divided into *θ*(*μm*)^3^ cubes. The midpoint of the lowest plane in the cube (u, v, z) was used as the three-dimensional geographic coordinates, and the number of cells of each type in the cube was recorded. The data in each of these cubes was used as input to the analysis and model.

Moran’s I index was used to measure spatial autocorrelation for each cell type^17^. The formula for Moran’s I index is

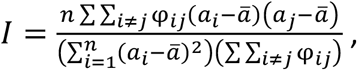

where *n* is the total number of cubes, a_i_ is the number of cells in a particular type at the *i*th cube, a_j_ is the number of cells at the *j*th cube, ā is the mean of the number of cells at each cube, and φ_ij_ is a spatial weight. The formula for φ_ij_ is

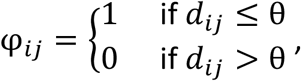

where d_ij_ be the Euclidean distance between the centroids of cube *i* and cube *j*.

The density-based spatial clustering of applications with noise (DBSCAN) algorithm was the algorithm used to cluster the cells^18^. For cube (u_i_, v_i_, z_i_), N_i_ = {(u_j_, v_j_, z_j_)|d_ij_ ≤ θ} is a set of all neighbouring cubes that are θ μm or less away from the *i*th cube. The number of cells in N_i_ is recorded as |N_i_|. When the faces of cubes are connected to each other, they are neighbours, so each cube in this case has six neighbours. If the total number of cells in cube (u_i_, v_i_, z_i_) and its neighbours is greater than 7, the number of cubes being considered, times γ, such as the third quartile of counts, then the *i*th cube and its neighbours are marked as high-density cubes. Let Ω = {(u_j_, v_j_, z_j_)||N_i_| ≥ 7γ, d_ij_ ≤ θ} is the set which includes all high-density cubes. For any cube(u_i_, v_i_, z_i_) ∈ Ω, *∂*_1_ = {(u_j_, v_j_, z_j_) |d((u_i_, v_i_, z_i_), Ω) ≤ θ} where *∂*_1_ includes all high-density cubes close to *i*th cube, and d(A, B) represents Euclidean distance between the set *A* and the set *B*. Then, *∂*_2_ = {(u_j_, v_j_, z_j_) |d(*∂*_1_, Ω) ≤ θ}, ⋯, *∂*_n+1_ = {(u_j_, v_j_, z_j_)|d(*∂*_n_, Ω) ≤ θ}. When |*∂*_n+1_| − |*∂*_n_| = 0 where |*∂*_n_| is the number of cubes in the *∂*_n_, iteration ends and *∂*_n_ is the first cluster which is records as C_1_. If Ω_1_ = Ω − C_1_ = ∅, then there is one cluster. Conversely, any cube (u_l_, v_l_, z_l_) ∈ Ω_1_ are selected. The second cluster C_2_ and Ω_2_ can be obtained using the same step. When Ω_p+1_ = ∅, p ∈ N^*^, the data has *p* clusters. In addition, for any cube that does not belong to any cluster, these cubes are in the set C_0_.

Permutation tests were used to detect changes in the number of cells in the cluster as well as the number of cells around the cluster^19^. The null hypothesis for the permutation test is that the mean number of cells in the cubes is independent of whether the cubes in C_τ_ or 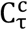. Here,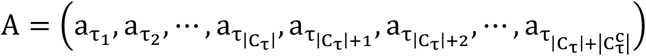 is an ordered observations set, where 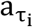 is the number of cells in the *i*th cube in the C_τ_. In the set *A*, the first |C_τ_| elements are the number of cells in the τth cluster, and the last 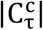 elements are the number of cells around the cluster. Hence, the real-valued statistic is used under this null hypothesis.

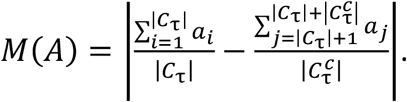

Then, a permutation π is created, that can reassign labels to individual datum. A reordered observation set is obtained:

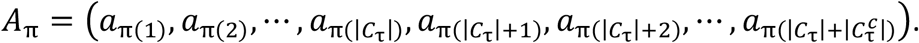

Similarly, a new reordered set also generates statistics M(A_π_). A collection of permutations Q can be characterised so that reordering {A_π_}_π∈Q_ are equally likely under the null hypothesis. Then, the empirical distribution of M(A_π_)_π∈Q_ is used to compare with M(A). Therefore, the Monte Carlo approximation with 500,000 test statistics was applied to estimate the two-tailed p-value, and the Holm-Bonferroni correction was performed.

The generalized linear model was used to predict the likelihood of the presence of cell types in a cluster versus cells not in a cluster. Each cube was changed into a dummy variable as the response variable based on whether it was in the cluster or not. The explanatory variables were binary variables, also determined by the presence or absence of the cell type. The generalized linear model was performed by ‘glm’ function in R (version 4.4.0).

## Supporting information

Supplemental Figure 1 wet protocol optimisation

## Code availability

All the submissions scripts used in this study can be found at: https://github.com/ga402/PACESS.

## ACKNOWLEDGMENTS

We thank staff of the core facilities at Imperial college London (CBS facility, FILM) and the Crick Institute (BRF facility) for their valuable help. This work was supported by a Royal Society/Irish Royal Society International Exchange Program IEC\R1\180061 to CLC and KRD; CRUK Programme Foundation Award C36195/A26770, Wellcome Trust Investigator Award 212304/Z/18/Z and a Fondation Alcea project grant to CLC; Science Foundation Ireland Project Grant 18/CRT/6049 to KRD. GA was funded by a Wellcome trust 4i clinical PhD studentship (203928/Z/16/Z) and an NIHR Academic Clinical Lectureship. We also acknowledge the National Institute for Health Research (NIHR) Biomedical Research Centre based at Imperial College Healthcare NHS Trust and Imperial College London. The opinions, findings and conclusions or recommendations expressed in this material are those of the author(s) and do not necessarily reflect the views of the NHS, the NIHR, the Department of Health or Science Foundation Ireland.

## FIGURES LEGEND

**Supplemental Figure 1:** Bone marrow thick section preparation.**(A)** Bone preparation workflow. Following harvest, the bones are cleaned, fixed (2 hours) and decalcified (10 days). Bones are embedded in 4% agarose and section on the vibratome at 250µm thickness. After heme removal, sections are incubated with specific antibodies and cleared before imaging. **(B)** Comparison of before vs after clearing of whole calvarium (left panels) and 250µm thick femur section (right panels). **(C)** Effect of clearing protocol on the bone marrow integrity. The same calvaria was first imaged in vivo by intravital microscopy before going through the staining/clearing protocol and re-imaged. The images show on the left part the IVM acquisition and on the right the post clearing results. Blood vessels (green) are identified by FITC dextran intravenous injection in the IVM image and endomucin (Endmcn) immunostaining on the processed bone. Mks are identified thanks to vWF reporter transgene (red). **(D)** Quantification of Mks’ size before (IVM) and after clearing protocol. **(E-F)** Effect of fixation time on immunostaining efficiency and background level using CD3 antibody (magenta). Representative images of each time point are shown in **(E)** and the ratio cell to background is plotted in **(F)**.

## REFERENCES

1. Friston, K.J. (2011). Functional and effective connectivity: a review. Brain Connect 1, 13–36. 10.1089/brain.2011.0008.

2. Mesa, K.R., Kawaguchi, K., Cockburn, K., Gonzalez, D., Boucher, J., Xin, T., Klein, A.M., and Greco, V. (2018). Homeostatic Epidermal Stem Cell Self-Renewal Is Driven by Local Differentiation. Cell Stem Cell 23, 677-686.e4. 10.1016/j.stem.2018.09.005.

3. Cheng, H., Zheng, Z., and Cheng, T. (2020). New paradigms on hematopoietic stem cell differentiation. Protein Cell 11, 34–44. 10.1007/s13238-019-0633-0.

4. Laurenti, E., and Göttgens, B. (2018). From haematopoietic stem cells to complex differentiation landscapes. Nature 553, 418–426. 10.1038/nature25022.

5. Kucinski, I., Campos, J., Barile, M., Severi, F., Bohin, N., Moreira, P.N., Allen, L., Lawson, H., Haltalli, M.L.R., Kinston, S.J., et al. (2024). A time- and single-cell-resolved model of murine bone marrow hematopoiesis. Cell Stem Cell 31, 244-259.e10. 10.1016/j.stem.2023.12.001.

6. Zhang, J., Wu, Q., Johnson, C.B., Pham, G., Kinder, J.M., Olsson, A., Slaughter, A., May, M., Weinhaus, B., D’Alessandro, A., et al. (2021). In situ mapping identifies distinct vascular niches for myelopoiesis. Nature 590, 457–462. 10.1038/s41586-021-03201-2.

7. Coutu, D.L., Kokkaliaris, K.D., Kunz, L., and Schroeder, T. (2018). Multicolor quantitative confocal imaging cytometry. Nat Methods 15, 39–46. 10.1038/nmeth.4503.

8. Nombela-Arrieta, C., Pivarnik, G., Winkel, B., Canty, K.J., Harley, B., Mahoney, J.E., Park, S.-Y., Lu, J., Protopopov, A., and Silberstein, L.E. (2013). Quantitative imaging of haematopoietic stem and progenitor cell localization and hypoxic status in the bone marrow microenvironment. Nat Cell Biol 15, 533–543. 10.1038/ncb2730.

9. Akinduro, O., Weber, T.S., Ang, H., Haltalli, M.L.R., Ruivo, N., Duarte, D., Rashidi, N.M., Hawkins, E.D., Duffy, K.R., and Lo Celso, C. (2018). Proliferation dynamics of acute myeloid leukaemia and haematopoietic progenitors competing for bone marrow space. Nature Communications 9, 519. 10.1038/s41467-017-02376-5.

10. Pirillo, C., Birch, F., Tissot, F.S., Anton, S.G., Haltalli, M., Tini, V., Kong, I., Piot, C., Partridge, B., Pospori, C., et al. (2022). Metalloproteinase inhibition reduces AML growth, prevents stem cell loss, and improves chemotherapy effectiveness. Blood Advances 6, 3126–3141. 10.1182/bloodadvances.2021004321.

11. Susaki, E.A., Tainaka, K., Perrin, D., Yukinaga, H., Kuno, A., and Ueda, H.R. (2015). Advanced CUBIC protocols for whole-brain and whole-body clearing and imaging. Nat Protoc 10, 1709–1727. 10.1038/nprot.2015.085.

12. Li, W., Germain, R.N., and Gerner, M.Y. (2017). Multiplex, quantitative cellular analysis in large tissue volumes with clearing-enhanced 3D microscopy (Ce3D). Proceedings of the National Academy of Sciences 114, E7321–E7330. 10.1073/pnas.1708981114.

13. Krivtsov, A.V., Twomey, D., Feng, Z., Stubbs, M.C., Wang, Y., Faber, J., Levine, J.E., Wang, J., Hahn, W.C., Gilliland, D.G., et al. (2006). Transformation from committed progenitor to leukaemia stem cell initiated by MLL-AF9. Nature 442, 818–822. 10.1038/nature04980.

14. Duarte, D., Hawkins, E.D., Akinduro, O., Ang, H., Filippo, K.D., Kong, I.Y., Haltalli, M., Ruivo, N., Straszkowski, L., Vervoort, S.J., et al. (2018). Inhibition of Endosteal Vascular Niche Remodeling Rescues Hematopoietic Stem Cell Loss in AML. Cell Stem Cell 22, 64-77.e6. 10.1016/j.stem.2017.11.006.

15. Aloysius, N., and Geetha, M. (2017). A review on deep convolutional neural networks. 2017 International Conference on Communication and Signal Processing (ICCSP), 0588–0592. 10.1109/ICCSP.2017.8286426.

16. Bodla, N., Singh, B., Chellappa, R., and Davis, L.S. (2017). Sof-NMS - Improving Object Detection with One Line of Code. In Proceedings - 2017 IEEE International Conference on Computer Vision, ICCV 2017 (Institute of Electrical and Electronics Engineers Inc.), pp. 5562–5570. 10.1109/ICCV.2017.593.

17. Espindola, G., Camara, G., Reis, I., Bins, L., and Monteiro, M. (2006). Parameter selection for region-growing image segmentation algorithms using spatial autocorrelation. International Journal of Remote Sensing 27, 3035–3040. 10.1080/01431160600617194.

18. Schubert, E., Sander, J., Ester, M., Kriegel, H.P., and Xu, X. (2017). DBSCAN Revisited, Revisited: Why and How You Should (Still) Use DBSCAN. ACM Trans. Database Syst. 42, 19:1-19:21. 10.1145/3068335.

19. Lehmann, E.L., and Romano, J.P. (2022). Testing Statistical Hypotheses (Springer International Publishing) 10.1007/978-3-030-70578-7.

20. Chanu, M.M., Lourembam, R., and Khelch, D. (2020). A Deep Learning Approach for Object Detection and Instance Segmentation using Mask RCNN. Journal of Advanced Research in Dynamic and Control Systems Volume 12, 95–104. 10.5373/JARDCS/V12SP3/20201242.

21. Lin, T.-Y., Dollár, P., Girshick, R., He, K., Hariharan, B., and Belongie, S. (2017). Feature Pyramid Networks for Object Detection. In 2017 IEEE Conference on Computer Vision and Pattern Recognition (CVPR), pp. 936–944. 10.1109/CVPR.2017.106.

22. Hawkins, E.D., Duarte, D., Akinduro, O., Khorshed, R.A., Passaro, D., Nowicka, M., Straszkowski, L., Scott, M.K., Rothery, S., Ruivo, N., et al. (2016). T-cell acute leukaemia exhibits dynamic interactions with bone marrow microenvironments. Nature 538, 518–522. 10.1038/nature19801.

23. Tissot, F.S., Gonzalez-Anton, S., and Lo Celso, C. (2024). Intravital Microscopy to Study the Effect of Matrix Metalloproteinase Inhibition on Acute Myeloid Leukemia Cell Migration in the Bone Marrow. Methods Mol Biol 2747, 211–227. 10.1007/978-1-0716-3589-6_17.

24. Shaifee, M.J., Chywl, B., Li, F., and Wong, A. (2017). Fast YOLO: A Fast You Only Look Once System for Real-time Embedded Object Detection in Video. J. Comp. Vis. Imag. Sys. 3. 10.15353/vsnl.v3i1.171.

25. Kharchenko, V., and Chyrka, I. (2018). Detection of Airplanes on the Ground Using YOLO Neural Network. 2018 IEEE 17th International Conference on Mathematical Methods in Electromagnetic Theory (MMET), 294–297. 10.1109/MMET.2018.8460392.

26. Redmon, J., Divvala, S., Girshick, R., and Farhadi, A. (2016). You Only Look Once: Unified, Real-Time Object Detection. In 2016 IEEE Conference on Computer Vision and Pattern Recognition (CVPR), pp. 779–788. 10.1109/CVPR.2016.91.

