## Supplementary figures and images for "PACESS: Practical AI-based Cell Extraction and Spatial Statistics for large 3D bone marrow tissue images"

### Supplemental Figure 1 wet protocol optimisation

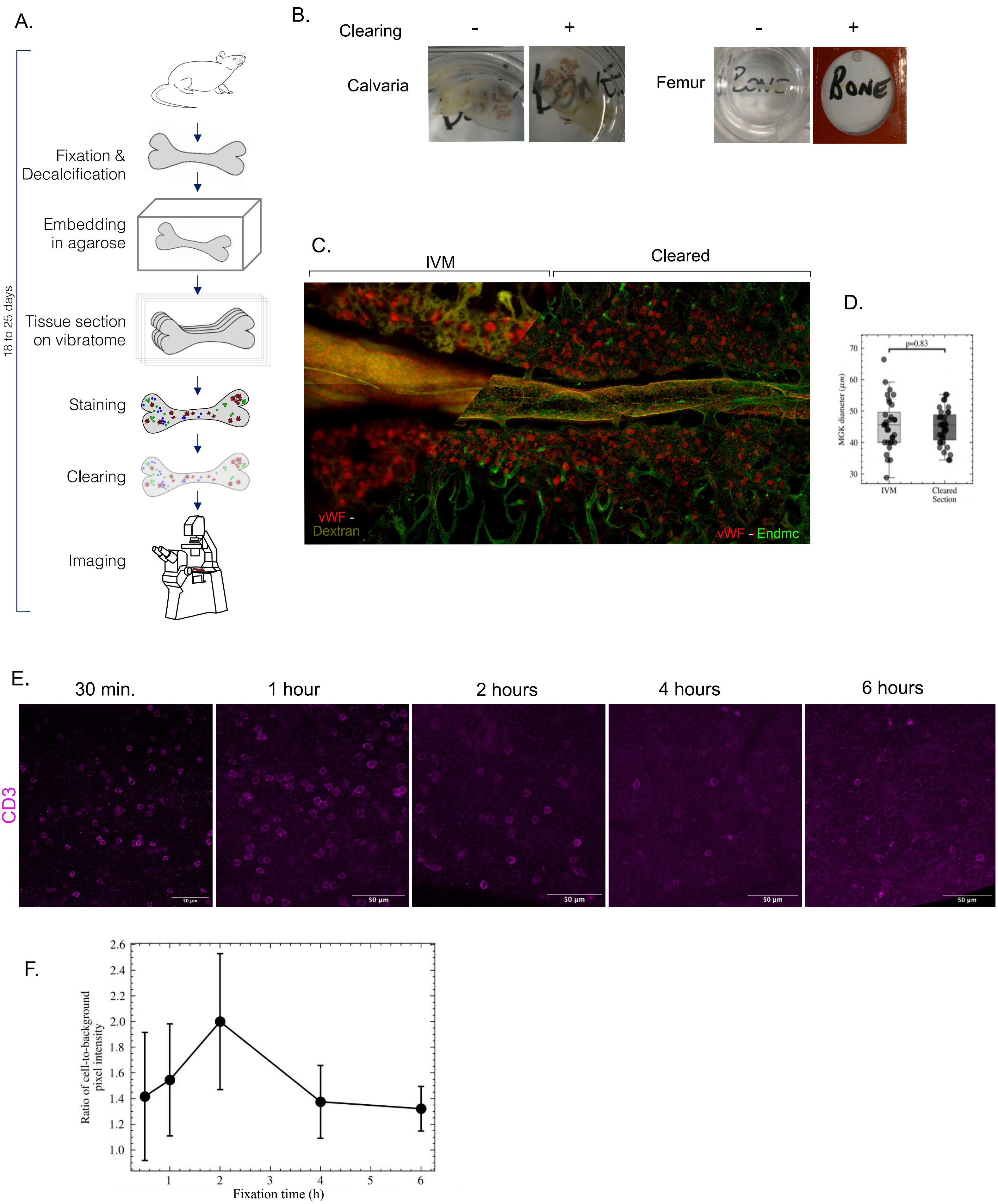
